# Increased EV yield and bioactivity of matrix-bound nanovesicles compared to conditioned culture medium-derived extracellular vesicles

**DOI:** 10.64898/2026.10.21.760728

**Authors:** Andrew White, Catherine Zhu, Vidya Sundaram, Hannah Levin, Marley J. Dewey

## Abstract

**Aim:** Extracellular vesicles (EVs) are nanoparticles secreted by cells for cell-cell communication functions, such as wound repair, but low yields often limit their use. Several isolation methods have been explored to increase EV yield from conditioned culture medium (liquid-EVs); however, few studies compare isolation techniques and yield between liquid-EVs and matrix-bound nanovesicles (MBVs), a novel EV subtype embedded within the extracellular matrix.

**Methods:** We isolate liquid-EVs and MBVs from mesenchymal stem cells (MSC) using size exclusion chromatography (SEC) and ultracentrifugation (UC) across various timepoints (14, 21, 24, 30 days). EVs are characterized by their yield, size, morphology, protein concentration, surface marker expression (CD9, CD81, ALIX), zeta potential, and influence on fibroblast wound closure.

**Results:** We find that isolation method influences yield, where both SEC isolated liquid-EVs and MBVs yield higher particle counts and protein compared to UC. Interestingly, we find that regardless of isolation method, MBVs have a significantly higher yield than liquid-EVs from the same culture (15.6-fold higher using UC and 7.49-fold higher using SEC). Culturing for 24 days yielded the greatest number of MBV particles and protein. We performed a fibroblast scratch assay to assess bioactivity and found that liquid-EVs decreased wound closure compared to a medium control, whereas MBVs did not.

**Conclusion:** Our work suggests SEC maximizes particle and protein yields for MSC-EVs. By achieving much higher yields of MBVs compared to liquid-EVs, and their promise as a wound repair strategy, we offer an alternative EV choice when EV yields are an issue towards translation.

## 1. Introduction

Extracellular vesicles (EVs) are lipid bilayer-enclosed nanoparticles secreted naturally by cells into the surrounding environment, where they can carry signaling molecules and elicit biological responses between cells^1^. Due to their cell-cell signaling nature, EVs represent promising cell-free therapies for tissue repair^2^. EVs have been isolated from a number of cell sources (mesenchymal stem cells, immune cells, blood cells, neural cells, others) with pronounced roles as potent immunomodulators for regenerative medicine applications, drug delivery vehicles for disease treatment, as well as diagnostic cell disease biomarkers^3^. Of particular interest are EVs isolated from mesenchymal stem cells (MSCs), as MSCs themselves are key cellular sources used in regenerative medicine applications ^4,5^. For example, EVs from human placenta-derived mesenchymal stem cells have been shown to accelerate collagen deposition and enhance wound healing in diabetic mouse models^6^. This is due to MSCs’ innate ability to undergo cellular reprogramming and differentiation, and paracrine signaling^7^. EVs are a major component of these paracrine factors, and interestingly, researchers have discovered that the therapeutic effects of MSCs could be largely driven by EVs and other paracrine factors^8^. For instance, MSC treatment in a rat osteoarthritis model had a negative effect on joint repair, whereas MSC-EV treatment had a more promising reparative effect by reducing inflammation and cartilage degeneration^9^. In another study, EVs derived from human bone marrow MSCs reduced liver fibrosis more effectively than the MSCs themselves, improving liver regeneration and reducing inflammation^10^. MSC liquid-EVs can also improve the rate of skeletal muscle regeneration, attenuate mitochondrial damage and inflammation, and reduce cell senescence^11^. These are just a few examples of MSC-EVs as therapies for tissue repair and regeneration.

Most MSC-based EV therapies used for tissue regeneration include EVs isolated from conditioned cell culture medium (liquid-EVs), often referred to as exosomes^12^. Liquid-EVs are free-floating EVs isolated from liquid sources such as blood, serum, conditioned culture medium, urine, among others, and the Minimal Information for Studies of Extracellular Vesicles (MISEV) defines exosomes as lipid bilayer vesicles smaller than 200 nm that are released from within the cell by the multivesicular body^13^. Previous research has strongly supported that liquid-EVs play a critical role in cell-cell communication, cargo delivery, cell proliferation, and angiogenesis, supporting their numerous benefits within regenerative medicine^14,15^. In recent years, a novel subtype of EV was discovered tightly embedded within the extracellular matrix (ECM), known as matrix-bound nanovesicles (MBVs)^16^. While numerous characteristics about liquid-EVs have been identified, comparatively little is known about MBVs. For example, liquid-EVs are formed via the Endosomal Sorting Complex Required for Transport (ESCRT) when mature endosomes form vesicles that are released from the cell, while MBV biogenesis remains unknown^17^. Nonetheless, numerous studies have highlighted differences between MBV and liquid-EV cargo, origin, and characteristics^18,19^. Specifically, it has been shown that MBVs have a unique phospholipid and microRNA cargo profile that differs from liquid-EVs^20^. While MBVs have demonstrated distinct differences from liquid-EVs through several studies^21^, they are also promising therapies for wound repair and immunomodulation. For example, MBVs administered both locally and systemically in rats reduced rheumatoid arthritis damage to bone tissue and joints^22^ and could also reduce bone resorption in an osteolysis mouse model^23^. Additionally, MBVs isolated from porcine urinary bladder matrix showed no cytotoxic response even in high doses (1×10^11^ particles/mL) compared to media-only controls^24^. These pro-regenerative effects of MBVs are thought to act through immunomodulatory pathways^22,,25–29^, including modulating the immune system through myeloid lineage cells^25^.

MSC-derived EVs offer an exciting alternative to cell-based therapies; however, low EV yields pose a major limitation, especially when translational use requires upscaling of EV production, which may impact both yield and cargo^30^. Although MSC MBVs could be a promising therapeutic alternative to liquid-EVs, we currently lack a comprehensive understanding of yield differences between these two EV subtypes and limited information on their comparative bioactivity. It is well recognized that EV source can also influence yield, as EVs isolated from conditioned media from skeletal muscle ex vivo tissue were 100 times greater than white adipose tissue EVs^31^. In addition, a chosen isolation method influences EV yield, with many methods, including ultracentrifugation, ultrafiltration, precipitation, size exclusion chromatography, among others, influencing this yield^32^. We therefore hypothesized that MBVs and liquid-EVs had significantly different yields, due to these EVs being from different sources (cell culture medium compared to ECM), even from the same cell population. We compared yields using ultracentrifugation (UC) and size-exclusion chromatography (SEC), as they are two of the most commonly used isolation techniques for EV^33^. Although many of these different isolation methods have previously been used to explore MBV yield^34^, these studies have yet to compare the yields of MBVs directly to liquid-EVs, and likewise few studies compare the bioactivity of these two EV populations side-by-side.

We aim to evaluate how various parameters (culture time period, isolation method) influence MBV and liquid-EV yield, characteristics, and bioactivity. Optimizing and comparing isolation outcomes of MSC MBVs and liquid-EVs allows us to address one of the major limitations in MSC-EV therapies: limited EV yield.

## 2. Materials and Methods

### 2.1. Mesenchymal stem cell culture and extracellular matrix sheet formation

Human bone marrow MSCs (RoosterBio, Male, 19 years old) at passage 6 were cultured on 150 mm petri dishes (2,222 cells/cm^2^) with 30 mL of exosome-depleted culture medium. Cell culture medium used in our studies contained 89% low-glucose DMEM (Fisher, Cat#11885092), 10% fetal bovine serum (FBS) (Genesee, Cat#25-514) depleted of exosomes, 1% antibiotic-antimycotic (Fisher, Cat#15240-062), and 50 µM L-ascorbic acid (TCI, Cat#A2521). To deplete our FBS of exosomes, we ultracentrifuged (Himac CP 80NX) FBS at 100,000 x g for 18 hours and collected the supernatant. MSC cultures were incubated at 37°C in 5% CO_2_. Medium was replaced every 3 days until EV isolation.

### 2.2. Liquid-EV isolation using ultracentrifugation

We collected conditioned cell culture medium every 3 days throughout the entire 21 day culture period or from only the last 3 days of culture, and both were immediately frozen at −80°C. After thawing the collected media, these were centrifuged at 500 x g for 10 min, 2,500 x g for 20 min, and 10,000 x g for 30 minutes, with the supernatant collected and the pellet discarded after each spin. Then, the remaining supernatant was filtered through a 0.22 μm filter and ultracentrifuged at 100,000 x g for 70 min. The resulting liquid-EV pellets were resuspended in filtered 1X PBS, aliquoted, and stored at −80°C for downstream applications.

### 2.3. MBV Isolation by ultracentrifugation

To isolate MBVs using ultracentrifugation, the cell culture medium was first removed (see Section 2.2 for cell culture medium processing), and the extracellular matrix (ECM) was washed twice with 10 mL of sterile PBS. Then, 12 mL of filtered Liberase buffer solution containing 0.01 mg/mL Liberase (Sigma, Cat# 5401151001), 1 M Tris-HCl, 5 M NaCl, 1 M CaCl2, and Milli-Q water was added to the dish, and using a cell scraper, the ECM was scraped in the buffer solution before being collected into conical tubes. The ECM and Liberase solution were briefly vortexed and incubated on a tube rotator at 37°C for 3 hours. Following incubation, the solution underwent similar EV isolation steps, starting with differential centrifugation and then ultracentrifugation, as outlined in Section 2.2. The resulting MBV pellets were resuspended in filtered 1X PBS, aliquoted, and stored at −80 °C for downstream applications.

### 2.4. Liquid-EV and MBV isolation by size-exclusion chromatography

EV isolation from conditioned cell culture medium and ECM digest followed similar steps outlined in Section 2.2 and 2.3, respectively. Briefly, culture medium or ECM digest was centrifuged at 500 x g for 10 min, 2,500 x g for 20 min, and 10,000 x g for 30 minutes, with the supernatant collected and the pellet discarded after each spin. Then, the remaining supernatant was filtered through a 0.22 μm filter. All EV samples were then concentrated down to 1 mL total volume using 100,000 kDa concentration columns (Fisher, Cat#UFC910096). This 1 mL sample was then added to a SEC column, packed with 10 mL of Sepharose beads (Sigma, Cat#CL2B300, Cross-linked, 70-230 mesh, 60-200 μm bead diameter). Liquid-EV and MBV fractions were determined by collecting eluted fractions 1 through 10. For each fraction, the total particles and total protein were used to create total particle-to-protein ratios (**Supp. Table 1**). Desired fractions had the highest particles and lowest proteins, which were fractions 3-5 for Liquid-EVs and fractions 3-6 for MBVs. Desired fractions were pooled, aliquoted, and stored at −80°C for downstream analysis.

### 2.5. Particle size and concentration quantification

To measure EV size and concentration, we utilized a NanoSight NS300 (Malvern Panalytical). Prior to the sample being run through the system, the syringe line was extensivley flushed with Milli-Q water. EV samples were serially diluted up to 1:1000, and higher when necessary. Samples were injected with a syringe pump at speed 100 and samples were flushed with Milli-Q before each new sample was loaded. All samples were set to camera level 14 and detection threshold 5. Videos (n=3) at 45 seconds each were taken per sample and analyzed by Nanoparticle Tracking Analysis.

### 2.6. Protein quantification

EV protein concentration was quantified utilizing a microBCA assay (ThermoFisher, Scientific Cat#23235). EV samples and kit standards were prepared using filtered 1X PBS as diluent. Following kit instructions, a working reagent was added to every well, and the plate was incubated at 37°C for 2 hours before absorbance was read at 562 nm on a spectrophotometer (Tecan Spark).

### 2.7. Western blotting

5 µg total protein per sample was used for all Western Blots. To each sample, Laemmli buffer (10% β-mercaptoethanol and 90% Laemmli buffer) was added, and this was heated to 95°C for 5 minutes. Samples were then centrifuged at max speed for 1 minute before being added to 10% Mini-PROTEAN TGX Precast Protein Gels (BioRad, Cat# 4561034). The bands were separated in 1X Running Buffer (25 mM Tris, 190 mM Glycine, 0.1% SDS) at 50 V for 25 minutes, followed by 150 V until the bands reached the bottom of the gel. Gels were transferred in 1X Transfer Buffer (25 mM Tris, 190 mM Glycine, 20% methanol) to a PVDF Membrane (Fisher Scientific, Cat#IPVH00010) sandwiched between filter paper (BioRad, Cat# 1703966). The membrane transfer occurred overnight (30 A, 4°C). Membranes were blocked with 2.5% non-fat dry milk in TBST (20 mM Tris pH 7.5, 150 mM NaCl, 0.1% Tween-20) and labeled with 1:2000 CD9 (Proteintech, Cat#20597-1-AP), 1:500 CD63 (Proteintech, Cat#25682-1-AP), and 1:5000 ALIX (Proteintech, Cat#12422-1-AP). A dilution of 1:2000 of secondary antibody (Proteintech, Cat#SA00001-2) was used. Secondary attachment was visualized by chemiluminescence using SuperSignal™ Western Blot Substrate Bundle, Pico PLUS (ThermoFisher, Cat#A43840). Bands were imaged using the Bio-Rad ChemiDoc MP Imaging System.

### 2.8. Transmission electron microscopy

5 µl of EV sample was pipetted onto a TEM grid (Fisher Scientific, Cat#50-260-38) and incubated at room temperature for 1 minute. Excess sample was removed with Whatman filter paper (Cytiva, Cat#1001-0155) and 5 µL of a 2% uranyl acetate negative stain was immediately added and removed, followed by another 5 µL negative stain, which was incubated at room temperature for 1 minute before being removed with filter paper. Grids were imaged using a Transmission Electron Microscope (Thermo Scientific, Talos F200X).

### 2.9. Zeta potential

EV samples were diluted in Milli-Q water to approximately 1 x 10^7^ EV/mL, placed in a clear disposable zeta cell, and zeta potential (mV) was measured using a Zetasizer (Malvern, Nano ZS) by averaging 3 runs per sample and taking the average of each data set. Instrument temperature was set to 25°C and the attenuator was set to 11.

### 2.10. Fibroblast scratch assay

Human dermal fibroblasts (ATCC, Cat#PCS-201-012, female, 41, caucasian) were seeded at 10,000 cells/well in culture-insert 2 well 24 ibiTreat plates (Ibidi, Cat#80242) and incubated at 37°C in complete DMEM (89% low-glucose DMEM, 10% FBS, 1% anti-anti) at 5% CO2 until cells were adhered and a confluent monolayer was formed at the bottom of the well plate in each compartment of the inserted well plate. After monolayer formation, each insert was removed using sterile tweezers and 1 mL of media was used to wash any cell debris resulting from removing the inserts. Liquid-EV (8 µg/well) and MBV (8 µg/well) were added to exosome-depleted DMEM media (89% low-glucose DMEM, 10% exosome-depleted FBS, 1% anti-anti) to wells for 16 hours. One medium control was used containing no EV treatment, which was complete DMEM (89% low-glucose DMEM, 10% exosome-depleted FBS, 1% anti-anti). All wells were imaged at hour 16 with a brightfield microscope (Olympus IX81 Inverted Compound Microscope), and images were analyzed using ImageJ’s ‘wound healing size tool’ using the ‘wound_healing_size_tool’ plugin^35^. Any wells that could not be analyzed with the plugin were measured manually using ‘area measurement’ function in ImageJ.

### 2.11. Statistical analysis

All data was checked for assumptions of normality and equal variance prior to running statistical tests. Normality was assessed by the Shapiro-Wilk test and equal variance using a Brown– Forsythe test. If assumptions were met, a one-way ANOVA using a Tukey’s post-hoc analysis was performed for 3+ groups, or a t-test for 2 groups. If another test was used (lack of normality or equal variance), this was denoted in the figure caption. All graphs and statistical analyses were made through Prism (GraphPad Software, Inc). A value of p < 0.05 was considered statistically significant and p-values are reported on all graphs. All graphical data are presented as mean ± standard deviation.

## 3. Results

### 3.1. Liquid-EV and MBV follow expected MISEV guidelines for size, morphology, and protein expression

We first confirmed that we effectively isolated liquid-EVs and MBVs from MSCs by examining EV size, morphology, and common EV surface markers. Representative NTA data demonstrated a mean size >200 nm range, which aligns with the expected small EV size range outlined in MISEV (**Fig. 1B, Supp. Fig. 1**). TEM images demonstrate expected cup-shaped EV morphology (**Fig. 1B, Supp. Fig. 2**). Western blot analysis demonstrates expression of CD9, CD63, and ALIX in both liquid-EVs and MBVs (**Fig. 1 B**, **Supp. Fig. 3).**

**Figure 1.**
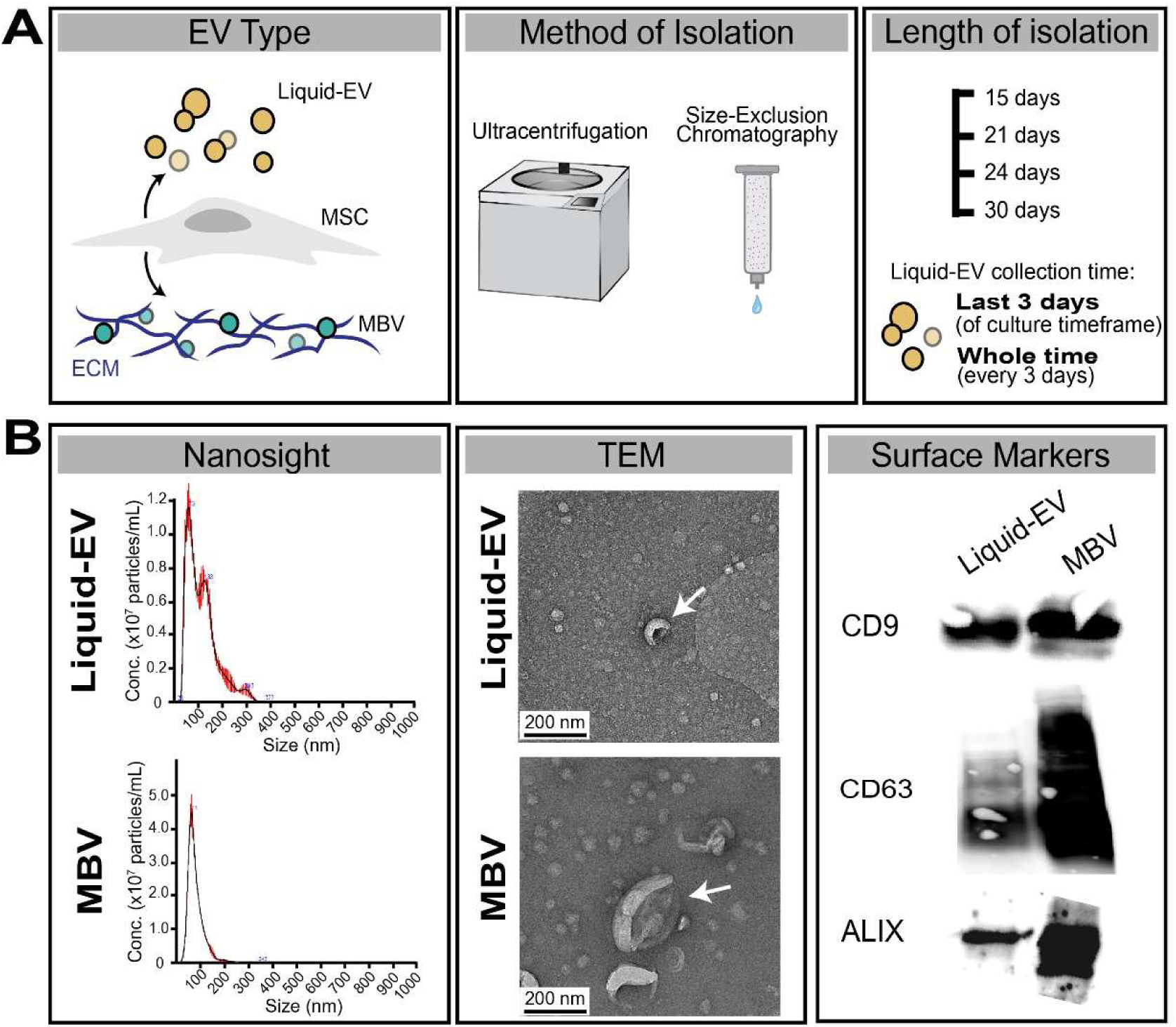
Experimental overview and characterization of MSC liquid-EVs and MBVs. (A) Schematic illustrating the two EV subtypes, methods of isolation, and the duration of performed within the study. (B) Representative data of Nanoparticle Tracking Analysis (NTA). Transmission Electron Microscopy (TEM), and western blots of MSC liquid-EVs and MBVs isolated after 21 days of culture.

### 3.2. Isolation methods alter EV particle and protein yields

We next examined how the isolation method, UC or SEC, influenced our resulting EV preparations. We achieved significantly higher total EV particles using SEC compared to UC regardless of EV type, liquid-EVs or MBVs (**Fig. 2A**). When isolating MBVs, SEC yielded (9.48 ± 0.58) x 10^11^ particles/mL compared to (5.56 ± 1.9) x 10^10^ using UC, representing a 17.1-fold increase in yield when using SEC. When isolating liquid-EVs the last 3 days of culture, SEC yielded a (1.27 ± 0.56) x 10^11^ compared to (3.56 ± 0.76) x 10^9^ using UC, representing a 35.5-fold increase in yield when using SEC. When isolating liquid-EVs across the entire culture period, SEC yielded (5.65 ± 1.70) x 10^10^ compared to (4.67 ± 0.53) x 10^9^ using UC, representing a 12.1-fold increase in yield when using SEC. Similarly, EVs isolated using SEC across all groups showed significantly higher levels of total protein compared to UC-isolated EVs. Likewise, SEC isolated EVs had greater protein yields of 8.48-fold for MBV, 6.73-fold for liquid-EVs (last 3 days), and 2.66-fold for liquid-EVs (whole time) compared to UC (**Fig. 2A**). This ultimately led to a higher particle-to-protein ratio in all EV groups using SEC compared to UC. While there was a significant difference in mean size of our MBVs isolated by SEC compared to UC, our liquid-EVs showed no significant difference. However, all groups were in the typical small EV size range (30-200 nm), which follows expected MISEV guidelines, as previously mentioned. Finally, the isolation method did not affect the zeta potential of any of our groups (**Fig. 2A**, **Supp. Fig. 4**).

**Figure 2.**
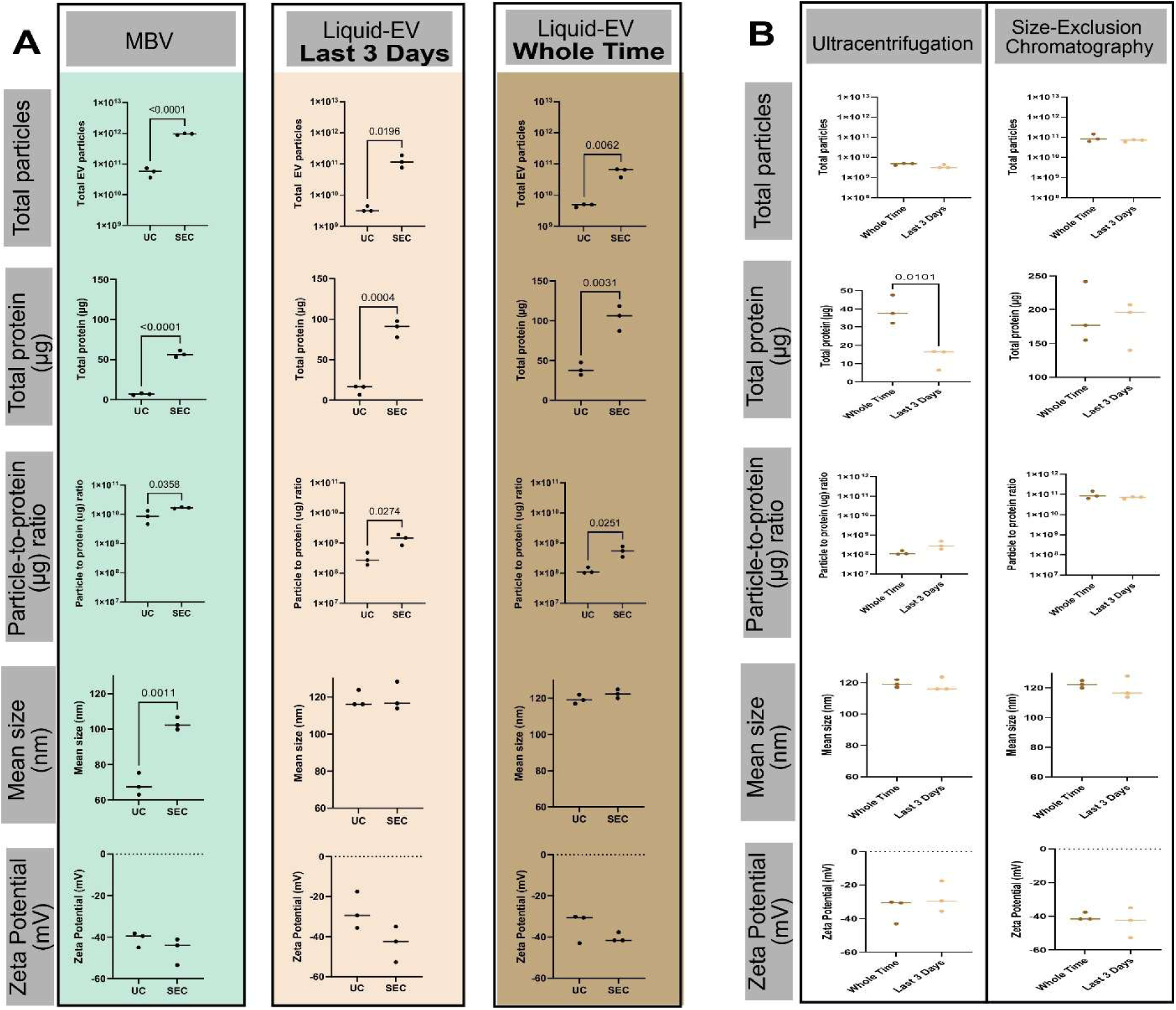
Comparison of EVs isolated using UC and SEC. (A) Graphs of UC compared to SEC for MBVs, and liquid-EVs collected from media during the last 3 days of the culture period (Last 3 days), and liquid-EV pooled from across the entire culture period (Whole Time, 21 days). Graphs are broken into categories of total particles, total protein, particle-to-protein ratios, mean size, and zeta potential. (B) Comparisons of liquid EVs isolated on the last 3 days of culture compared to the entire culture period when isolated by either UC or SEC. Similarly, groups were broken into the same categories as in (A). Data are expressed as mean ± standard deviation (n=3) and significant differences are represented as p-values. T-tests were performed for all groups.

### 3.3. Liquid-EV particle yield is not impacted when isolated during the entire culture period compared to the last 3 days of culture

We then wanted to determine if the timeframe of collecting our culture media influenced our liquid-EV yield. Therefore, we examined isolating EVs from conditioned culture medium either across the whole culture period of 21 days (Whole Time) or just the last 3 days of culture (Last 3 Days). Regardless of using either UC or SEC to isolate EVs, we found no significant difference in particle yield between our two liquid-EV conditions (**Fig. 2B**). This trend is consistent for the mean size and zeta potential of our groups. We did find an increase in total protein yield of liquid-EVs isolated from the Whole Time group using UC; however, this ultimately was not a significant difference for comparisons of particle-to-protein ratios between our groups.

### 3.4. MBV particle yields are greater compared to liquid-EVs regardless of isolation method

We sought to directly compare how isolation methods influence our liquid-EV and MBV total particle yields. We achieved significantly higher (p < 0.05) total MBV particles compared to liquid-EVs regardless of whether we utilized SEC or UC (**Fig. 3A, B**). UC yielded an average (5.56 ± 1.88) x 10^10^ MBVs compared to (3.56 ± 0.76) x 10^9^ liquid-EVs, representing a 15.62-fold increase in yield of MBVs compared to liquid-EVs. SEC also yielded an average (9.48 ± 0.58) x 10^11^ MBVs compared to (1.27 ± 0.56) x 10^11^ liquid-EVs, representing a 7.49-fold increase in yield of MBVs compared to liquid-EVs. In contrast, when isolating EVs using SEC, our liquid-EVs had significantly higher total protein compared to our MBVs. This method yielded 88.80 ± 10.13 µg total protein of liquid-EVs compared to 56.84 ± 4.16 µg total protein for MBV, representing a 1.56-fold increase in protein yield of liquid-EVs compared to MBVs (**Fig. 3B**). Finally, we show that MBVs have higher particle-to-protein ratios compared to liquid-EVs across both UC and SEC. For UC, MBVs had an average (8.76 ± 4.27) x 10^9^ particles/μg compared to liquid-EVs average of (1.51 ± 3.13) x 10^8^ particles/μg. For SEC, MBVs had an average (1.67 ± 0.10) x 10^10^ particles/μg compared to a liquid-EV average of (1.42 ± 0.54) x 10^9^ particles/μg. While there was a significant difference in total protein between our EV types when isolated by SEC, our groups showed no significant difference when isolated by UC (**Fig. 3A**).

**Figure 3.**
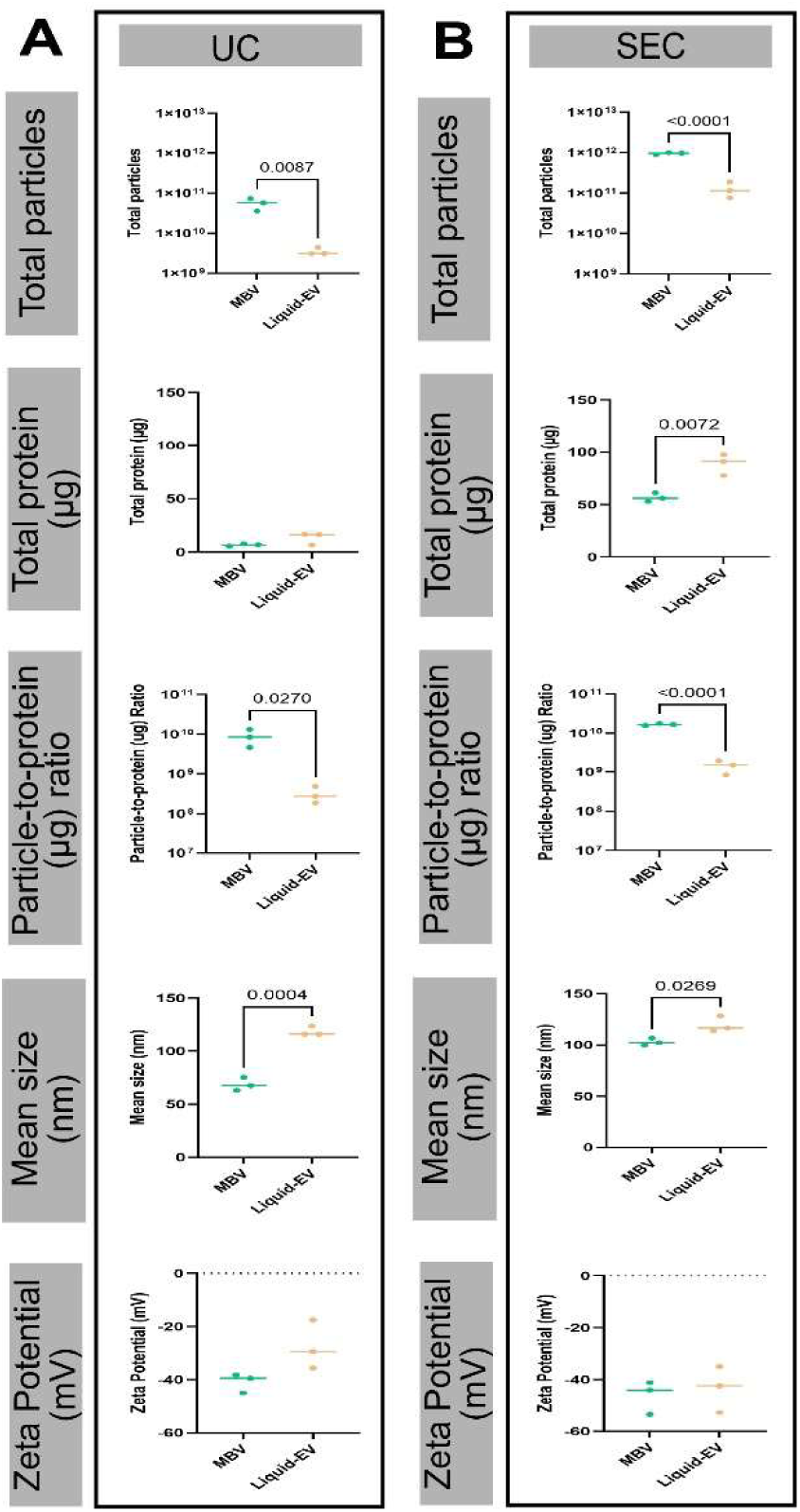
Comparison of MBVs and liquid-EVs isolated using UC and SEC. (A) Graphs of MBVs compared to liquid-EVs (last 3 days) when isolated by ultracentrifugation (UC). Graphs are broken into categories of total particles, total protein, total particle to protein ratios, mean size, and zeta potential. (B) Graphs of MBVs compared to liquid-EVs (last 3 days) when isolated by size exclusion chromatography (SEC). Similarly, groups were broken into the same categories as (A). Data are expressed as mean ± standard deviation (n=3) and significant differences are represented as p-values. T-tests were performed for all groups.

We also compared the average size and zeta potential between both liquid-EVs and MBVs when isolated using either method. We found a significantly larger mean size (p < 0.05) of our liquid-EVs compared to our MBVs regardless of isolation method (**Fig. 3A**). When isolated by UC, our MBV and liquid-EV sizes were 68.57 ± 6.23 nm and 121 ± 4.93 nm, respectively. Whereas, when isolated by SEC, our MBV and liquid-EV sizes were 103 ± 3.50 nm and 120 ± 7.65 nm, respectively. We see no difference in our zeta potential between the groups for either isolation method (**Fig. 3A**).

### 3.5. MBV particle count and protein concentration peak at 24 days of culture

We then sought to investigate any relationship between length of cell culture time and EV yield. Using SEC, we achieved the highest total EV particles and protein when isolating MBVs at 24 days of culture (**Fig. 4A**). For MBVs, we yielded (1.55 ± 0.39) x 10^12^ total particles at 24 days, compared to (5.93 ± 3.81) x 10^11^ total particles at 15 days, (9.48 ± 0.58) x 10^11^ total particles at 21 days, and (9.80 ± 6.74) x 10^11^ total particles at 30 days. Overall, 24 days represented a 2.61-fold increase from 15 days, a 1.63-fold increase in yield from 21 days, and a 1.58-fold increase in yield from 30 days. For liquid-EV, we yielded (1.13 ± 0.48) x 10^11^ total particles at 15 days, compared to (1.27 ± 0.57) x 10^11^ total particles at 21 days, (5.16 ± 3.17) x 10^10^ total particles at 24 days, and (2.95 ± 0.56) x 10^10^ total particles at 30 days. Overall, we see no significant difference in particle yield between any of the isolation time points for liquid-EVs (**Fig. 4B**). We observed a similar trend in protein concentration as we did with particle yields. The greatest protein yield for MBVs was at day 24, with a yield average of 92.91 ± 4.83 ug, while culture timeframe did not impact liquid-EV protein yield. We found no significant difference in protein-to-particle ratios across all days of either MBVs or liquid-EVs. Taken together, our data indicate that there is an optimal yield culture timeframe for MBVs at 24 days, but not liquid-EVs.

**Figure 4.**
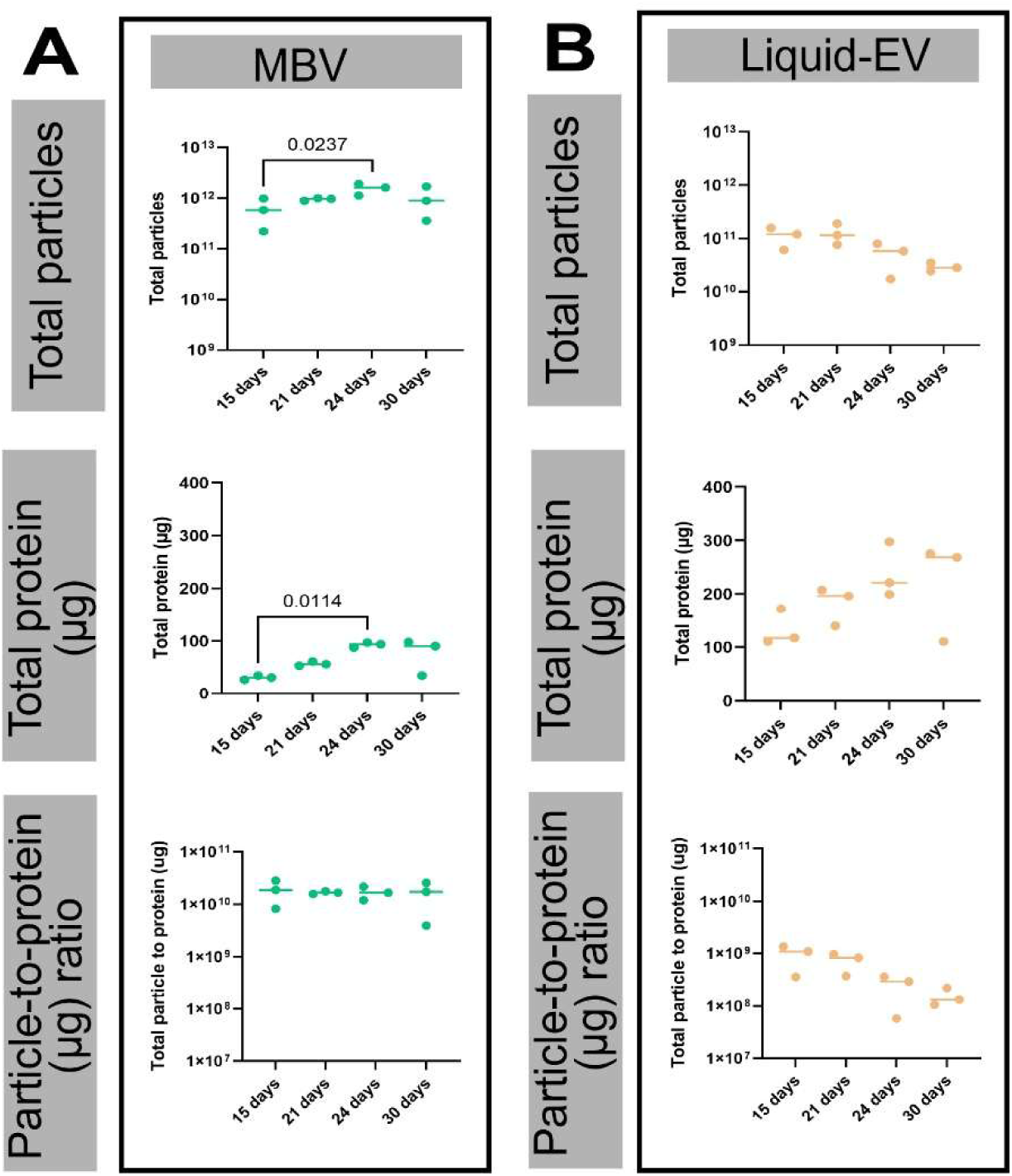
Comparison of MBVs and liquid-EVs across various culture periods. (A) Graphs of MBVs yield when isolated across various culture lengths (day 15, 21, 24, 30). Graphs are broken into categories of total particles, total protein, and particle-to-protein ratios. (B) Graphs of liquid-EV yields when isolated across various culture lengths (day 15, 21, 24, 30). Similarly, groups were broken into the same categories as (A). Data are expressed as mean ± standard deviation (n=3) and significant differences are represented as p-values. A One-way ANOVA was performed for all groups.

### 3.6. Liquid-EVs inhibit wound closure compared to a medium control

We assessed bioactivity differences between liquid-EVs and MBVs by measuring the percent closure of a scratch on a monolayer of fibroblasts after 16 hours with EV treatment. Compared to a medium control, liquid-EVs performed significantly (p < 0.05) worse, with less closure than the control (**Fig. 5A**). In contrast, MBVs had no significant difference in closure compared to the control; however, MBVs were also not significantly different from liquid-EVs. Overall, the average closure of the media control was 73.36 ± 6.00%, the liquid-EV group was 58.26 ± 8.40%, and the MBV group was 74.48 ± 16.76% (**Fig. 5B**). Of note, the MBV group average closure was higher than the control but had a very high standard deviation and variability between groups.

**Figure 5.**
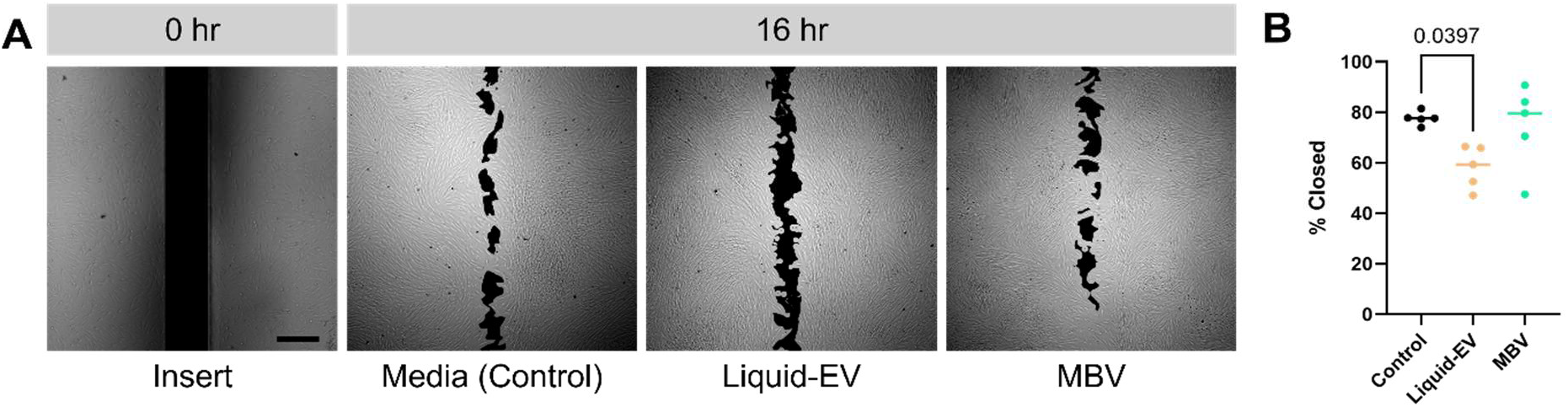
Human dermal fibroblast scratch assay with EV treatments. (A) Representative images (hour 0, hour 16) of insert, media control, Liquid-EV, MBV. Area analyzed is colored black for clarity. (B) Graph of percent closure between all conditions. Data are expressed as mean ± standard deviation (n=5) and significant differences are represented as p-values (p < 0.05). One-way ANOVA performed for all groups.

## 4. Discussion

Our study explored both the impact of SEC and UC isolation methods on the resulting yield of liquid-EVs and MBVs isolated from MSCs, as well as comparing characteristics and bioactivity of these two EV populations, to address an EV translational challenge related to low yields.

We first verified successful isolation and characteristics of MSC liquid-EVs and MBVs. Related to their size, MISEV states that small liquid-EVs remain below 200 nm, which was consistent with our findings. For MBV characteristics, such as size range and zeta potential, our findings were also consistent with published works, where average zeta potential has been reported to be −42.5 ± 3.42 mV for MSC MBV and −33.3 ± 2.54 mV for MSC liquid-EV, and average size was 117.8 ± 1.56 nm for MSC MBV and 162.1 ± 10.9 nm for MSC liquid-EV^21^, which is similar to our findings that MSC MBV have lower zeta potential and smaller sizes than liquid-EVs. Our MBVs and liquid-EVs were also enriched in CD9, CD63, and ALIX, common EV surface markers^21^.

One of the objectives of our study was to examine the difference between utilizing SEC or UC when isolating both liquid-EVs and MBVs. We found that EV yield is higher using SEC isolation than UC, regardless of whether MBVs or liquid-EVs were being isolated. This is in alignment with literature, as a study investigating umbilical cord blood mononuclear cell derived-EVs found that a combination of ultrafiltration and SEC had a yield 400 times higher than when isolated by traditional UC (per mL of media). In addition, this combined isolation method resulted in EVs with lower protein and lipid contamination than EVs isolated by UC, while the biologically relevant contents that promote bioactivity were retained^36^. Previous research has also examined UC and SEC independently, where rat plasma-derived EVs isolated by SEC had higher particle and protein amounts than when isolated by UC^37^. Another study isolated EVs from primary human cardiac mesenchymal-derived progenitor cells and found that SEC achieved higher particle concentrations than UC^38^. These studies all demonstrated that SEC achieves higher yields than UC for liquid-EVs, in alignment with our findings. Interestingly, in a study comparing MBV particle and protein yields across various isolation methods, authors found that MBV particle-to-protein ratios were similar between UC and SEC groups, which is different from our own findings of a higher particle-to-protein ratios using SEC than UC^39^. This could be due to the impact that cell source has on EV characteristics^40^. Other papers comparing UC and SEC for various types of EV have also suggested that SEC has similar or better performance compared to UC in terms of isolating pure and functional EVs^41^. For example, one study compared UC against SEC for isolation of liquid-EV (exosomes) from rat plasma and found that when isolated by SEC, there were higher particle number, protein content, particle/protein ratios, and signal of biomarkers compared to liquid-EV isolated by UC^37^. This is likely due to shear forces during UC, which can disrupt EV function and form, and aggregation of the EV particles during UC may lead to UC-isolated EVs appearing larger in size^42^. Due to this, we decided to focus our attention on using SEC-isolated EVs for later studies of timing and bioactivity assays.

We next investigated the impact of the timing of liquid-EV media collection. For SEC groups, we found that there was no difference in yield of liquid-EVs isolated from the last three days compared to EVs isolated from the entire 21-day culture period. For UC, we did see higher protein yields when isolating from the entire culture period compared to only the last three days; however, SEC yields overall for both particles and protein were still higher than UC groups. Therefore, we moved forward with collecting conditioned medium from the last three days of culture for our future experiments. There is minimal existing research on the impact of culture collection time on EV isolation for mammalian-derived EVs. However, others have examined the effect of storage conditions on EV yield. Other researchers have found that EV isolated from a plasma source that had been stored at −80 °C for 6 months showed decreased yield (particles/mL) compared to when isolated fresh or when stored at −80 °C for 4 weeks. They found a similar trend in EV yield when measured fresh, or after being stored at −80 °C for 4 weeks or 6 months^43^. These results could explain why we see no difference in liquid-EV yield after collecting medium for 3 days compared to the entire culture period, as our ‘whole time’ media conditions were stored at −80 °C throughout the culture period of 21 days. Alternatively, this could also be due to larger processing volumes of media, which could result in a higher loss of EVs due to the need for more concentration columns and concentration steps. For our whole time group, we processed 600% more medium than the last three days group.

We then directly compared the yield of MBVs to liquid-EVs and we found that the total yield of MBVs is higher than liquid-EVs, regardless of UC or SEC isolation. This aligns with available literature, where MSC-derived liquid-EVs showed an average of 2.095 ± 0.2 femtogram of protein per particle compared to MBVs, which showed 0.042 ± 0.006 femtogram per particle^21^. While we could not find direct yield comparisons between MBVs and liquid-EVs, other papers have compared their size differences. For example, researchers have previously noted similarities in size between liquid-EVs and MBVs derived from 3T3 fibroblast cell cultures^20^. This is different from our results, which have demonstrated a difference in size between MBVs and liquid-EVs. This could be due to cell origin and MBV processing conditions, as this study compared fibroblast EVs, including decellularized ECM, and we isolated MSC EVs from ECM that was not decellularized.

We then explored how EV yield changed across various isolation time points (15 days, 21 days, 24 days, 30 days). Our results suggest that the optimal time to culture MSCs to obtain the highest EV yield may differ depending on whether MBVs or liquid-EVs are being collected. While we did not observe significant differences for liquid-EVs, for MBVs we found that the ideal culture length may sit between 15 and 30 days, around 24 days. While there has never been an optimal culturing condition timeframe for MBVs reported previously, other researchers have cultured umbilical cord MSCs for 14 days to allow for ECM accumulation^45^.

Finally, our study measured differences in wound healing between liquid-EVs and MBVs using a fibroblast scratch assay. We see comparative bioactivity responses (% closure) between our MBV and media control group; however, our liquid-EV group had a significantly lower amount of closure than our media control, potentially inhibiting closure. This indicates that MSC MBV may have better bioactivity than liquid-EV in wound closure applications, as MBVs did not inhibit closure. Of note, MBV had a large range in bioactivity, with some samples actually increasing closure (91%) and others inhibiting (47%) compared to the medium control average (78%). In other studies, liquid-EVs and MBVs from multipotent mesenchymal stromal cells were applied to both a scratch assay and tubulogenesis assay, showing that both EV types enhance cell migration and that MBVs have higher angiogenic potential^44^. Researchers have also demonstrated that liquid-EVs may have a dose-dependent effect on wound closure for normal adult fibroblasts^45^. The potential reason we saw an inhibition of closure, rather than enhancement like other published works, could be due to culture conditions. For example, researchers have shown that lower seeding densities improved EV yield per cell and that utilizing trypsinization above passage 4 decreased bioactivity^46^. In our study, we isolated MSC EVs at passage 6, which could account for less optimal bioactivity according to this study.

From a clinical perspective, reported therapeutic doses of EV in clinical trials consist of high particle counts, requiring 10^8^ to 10^11^ particles per dose per patient^47,48^. Manufacturing these therapies at scale would require an efficient, optimized EV isolation process that minimizes possible product loss. From this work, we achieved a yield on the order of 10^11^ total MBVs from a single 150 mm petri dish, compared to on the order of 10^10^ liquid-EVs. This further demonstrates isolating MBVs may be easier to achieve required patient doses of EVs than liquid-EVs, and within achievable ranges for clinical applications. However, further *in vivo* results would be necessary to confirm MSC MBV are of similar, or better, bioactivity than liquid-EVs.

There are a few limitations to our study. Firstly, our study focused on yield and bioactivity of MSC-EVs, but did not explore other cell types, and as such, the yield and bioactivity between liquid-EVs and MBVs may differ depending on the cell type. There are also several appropriate EV isolation methods, and while our study compared two of the most common isolation methods (SEC and UC), other researchers have supported the benefit of combining multiple techniques to achieve optimal yields. Therefore, an even greater yield of MBVs could potentially be achieved with combinatorial methods. While there are also many bioactivity assessments, we focused on a scratch assay, and in the future we plan to examine the bioactivity differences between liquid-EVs and MBVs in a more comprehensive manner, including *in vivo* experiments. Finally, our study did not decellularize our ECM prior to MBV isolation. The effects of ECM decellularization on MBVs are currently unknown, as some detergents used in decellularization processes such as SDS and Triton-X can also damage EVs^49^. In the future, we plan to explore whether decellularization is a necessary step prior to MBV isolation, and if this alters MBV yield.

## 5. Conclusion

Our work highlights several key takeaways: SEC yields a significantly higher number of MSC EV particles than UC regardless of EV type (liquid-EV or MBV), and the culture timeframe of MSC MBV isolation impacts yield, with a ‘peak’ of 24 days to maximize efficiency of our isolations. Interestingly, we see significantly higher MSC MBV particle yields regardless of isolation method compared to liquid-EVs. Finally, our bioactivity results demonstrate that MSC liquid-EVs decreased wound closure compared to a medium control, whereas MBVs did not. Thus, MBVs may represent a bioactive and higher-yield source of MSC EVs for regenerative medicine applications than liquid-EVs.

## Acknowledgments

This study utilized instrumentation provided by the University of California – Santa Barbara’s Biological Nanostructures Laboratory, the Neuroscience Research Institute-Department of Molecular, Cellular and Developmental Biology Microscopy Facility, and the Materials Microscopy and Microanalysis Facility. We would also like to thank Dr. Arda Genc and Austin DuBose for assistance with TEM training, Dr. Jen Smith for Nanosight training, and other members of the Dewey Laboratory including Ivan Chavez, Max Mahoney-Schaefer, Haley Obenshain, and Mulunji Sampson for their feedback and assistance in this project.

## Authors’ Contributions

The following author contributions utilizes the Credit framework:

**Andrew White** – conceptualization, data curation, formal analysis, investigation, methodology, supervision, validation, visualization, writing − original draft, writing – review and editing.

**Catherine Zhu** – conceptualization, formal analysis, investigation, visualization, writing – original draft, writing – review and editing

**Vidya Sundaram** − conceptualization, formal analysis, investigation, visualization, writing – original draft, writing – review and editing

**Hannah Levin** – formal analysis, investigation, visualization, writing – original draft, writing – review and editing

**Marley Dewey** – conceptualization, funding acquisition, project administration, supervision, writing-review and editing

## Availability of Data and Materials

Data will be made available upon reasonable request by the authors.

## AI and AI-assisted tools Statement

Not applicable.

## Financial Support and Sponsorship

This work was partially supported by the Hellman Family Foundation Faculty Fellowship (MJD) and the National Institutes of Health NIGMS (R35GM160216) (MJD). This work was also partially supported by a National Institutes of Health T32 Training Grant (1T32GM141846) (AW) as well as the California Institute for Regenerative Medicine (CIRM) COMPASS program (CZ). The content is solely the responsibility of the authors and does not necessarily represent the official views of the National Institutes of Health.

## Notes

### Competing Interest Statement

The authors have declared no competing interest.

