## Supplementary files for "Increased EV yield and bioactivity of matrix-bound nanovesicles compared to conditioned culture medium-derived extracellular vesicles"

**Supplementary Figures**

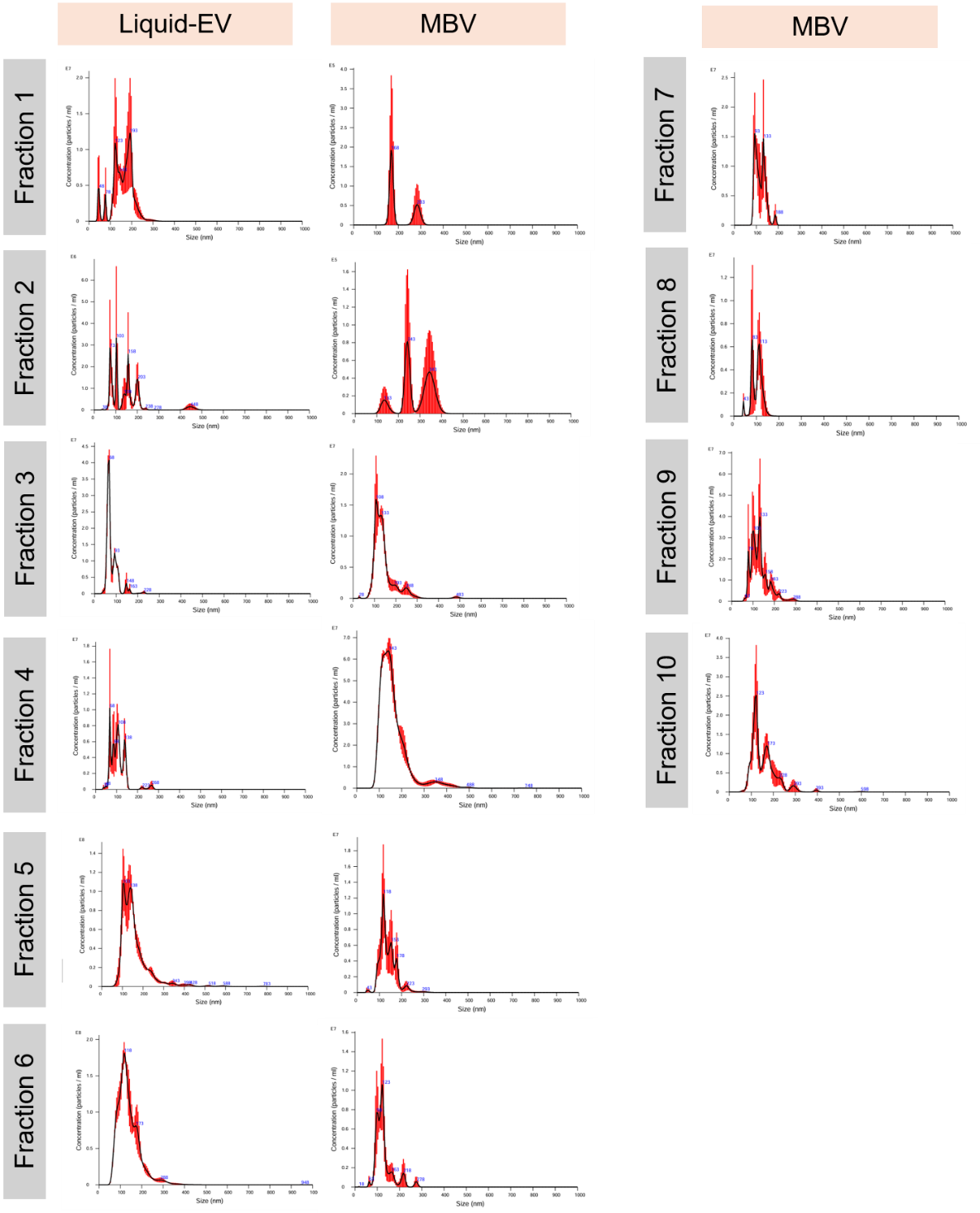

**Supp. Fig. 1. MBV and liquid-EV SEC fraction NanoSight plots**. Images of NanoSight plots of each of the collected fractions 1-10 for MBV and 1-6 for liquid-EV. Liquid-EV NanoSight plots were not collected for fractions 7-10 due to overwhelming protein numbers and low purity of EV.

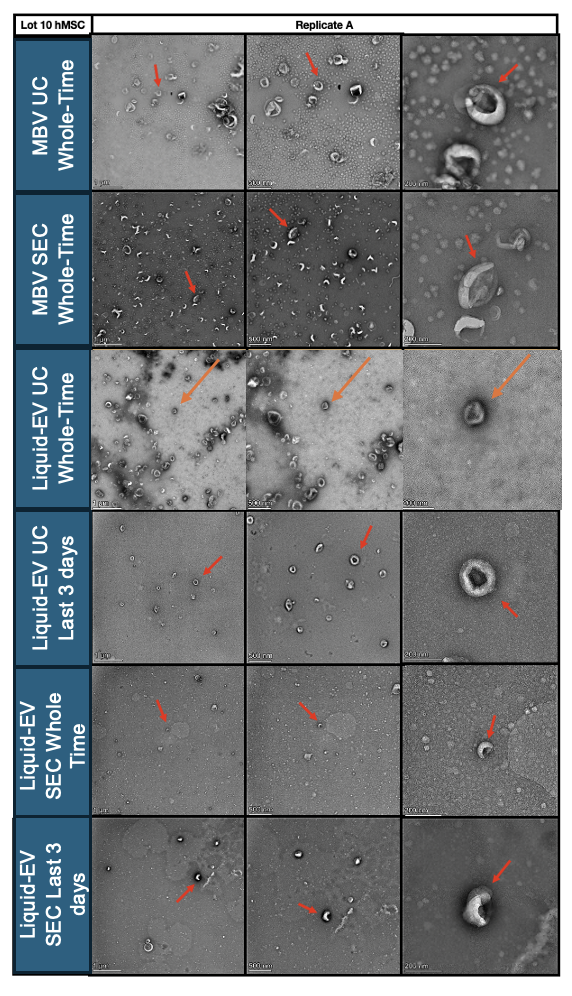

**Supp. Fig. 2. MBV and liquid-EV Transmission Electron Microscopy Results**. Images of NanoSight plots of each of the groups, which includes liquid-EV isolated from UC (whole culture period, last three days) or SEC (whole culture period, last three days), and MBV isolated by UC or SEC. Arrows are pointing to individual EV and scale bars are shown.

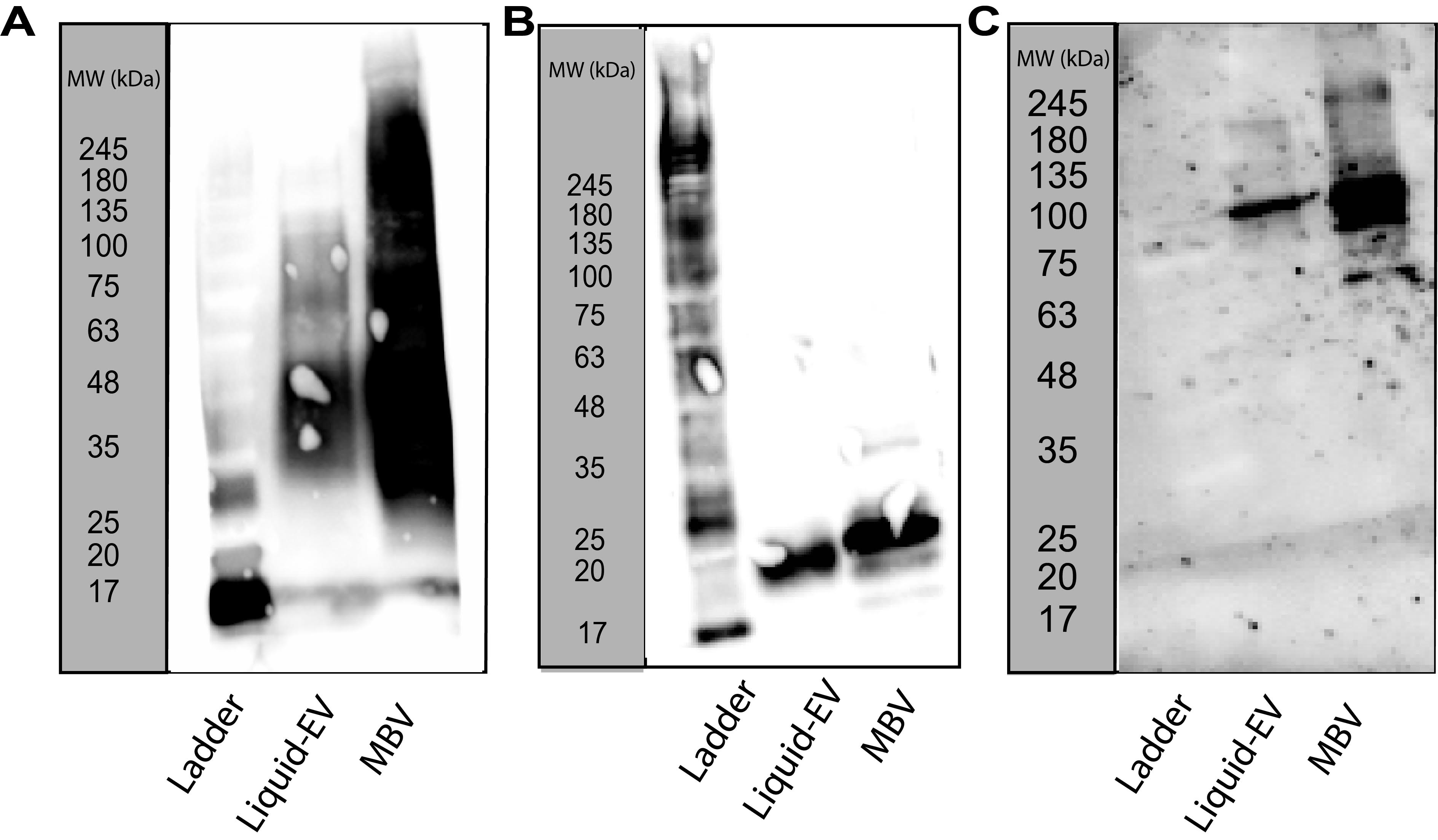

**Supp. Fig. 3. MBV and liquid-EV Western blots**. Images of WB gels of both liquid-EV and MBV for (A) CD63, (B) CD9, and (C) ALIX. Ladder is labeled with molecular weights.

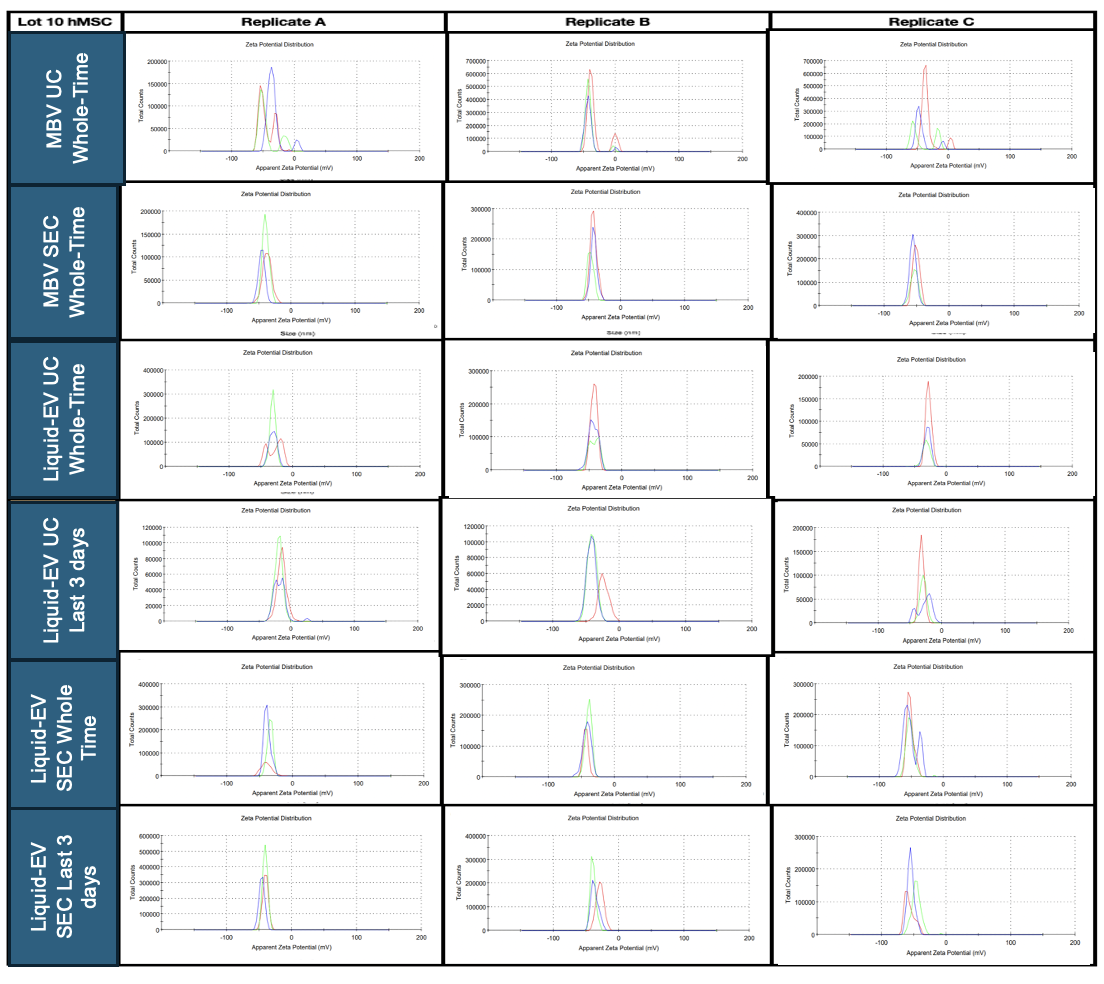

**Supp. Fig. 4. MBV and liquid-EV Zetasizer plots**. Images of zetasizer plots of each of the groups, which includes liquid-EV isolated from UC (whole culture period, last three days) or SEC (whole culture period, last three days), and MBV isolated by UC or SEC.

**Supplementary Tables**

**Supp. Table 1. SEC fraction determination for MSC liquid-EVs and MBVs.** Green outlines fractions used to isolate liquid-EV or MBV. “OVER” represents readings that were beyond the standard curve and represented protein concentrations over 200 µg/mL. N/A represents samples not collected due to extremely high protein concentrations.

| **Sample** | **Fraction** | **Total particles** | **Total Protein Concentration (µg)** | **Particles/Protein** |
| --- | --- | --- | --- | --- |
| MBV | 1 | 1.16E+07 | -0.92 | -1.27E+07 |
| MBV | 2 | 1.19E+06 | -0.83 | -1.43E+06 |
| MBV | 3 | 2.04E+11 | 2.30 | 8.87E+10 |
| MBV | 4 | 1.29E+12 | 12.85 | 1.00E+11 |
| MBV | 5 | 1.25E+11 | 4.94 | 2.53E+10 |
| MBV | 6 | 9.84E+10 | 4.40 | 2.24E+10 |
| MBV | 7 | 1.41E+10 | 13.26 | 1.06E+09 |
| MBV | 8 | 5.09E+09 | 45.32 | 1.12E+08 |
| MBV | 9 | 4.86E+09 | 62.18 | 7.82E+07 |
| MBV | 10 | 3.12E+09 | 44.00 | 7.09E+07 |
| Liquid-EV | 1 | 1.94E+08 | 1.6573725 | 1.17E+08 |
| Liquid-EV | 2 | 3.02E+07 | 0.77994 | 3.87E+07 |
| Liquid-EV | 3 | 2.11E+11 | 5.5570725 | 3.80E+10 |
| Liquid-EV | 4 | 9.36E+11 | 24.0156525 | 3.90E+10 |
| Liquid-EV | 5 | 1.97E+09 | 53.490885 | 3.68E+07 |
| Liquid-EV | 6 | 3.18E+09 | >200 (OVER) | N/A |
| Liquid-EV | 7 | N/A | >200 (OVER) | N/A |
| Liquid-EV | 8 | N/A | >200 (OVER) | N/A |
| Liquid-EV | 9 | N/A | >200 (OVER) | N/A |
| Liquid-EV | 10 | N/A | >200 (OVER) | N/A |
